# A mathematical description of non-self for biallelic genetic systems in pregnancy, transfusion, and transplantation

**DOI:** 10.1101/2024.04.10.588831

**Authors:** Klaus Rieneck

**Author notes:** Corresponding author: Klaus Rieneck, Department of Clinical Immunology, Laboratory of Blood Genetics, Rigshospitalet, Copenhagen University Hospital, Blegdamsvej 9, DK-2100, Copenhagen, Denmark. Transplantation; STR: Short Tandem Repeat. The author has no relevant financial or non-financial competing interests to disclose.

## Abstract

A central issue in immunology is the immunological response against non-self. The prerequisite for a specific immunological response is the exposure to the immune system of a non-self-antigen. Mathematical equations are presented, that define the fraction of all outcomes with a non-self-allele in biallelic systems at the population level in pregnancy and transfusion/transplantation medicine. When designing assays, the mathematical descriptions can be used for estimating the number of genetic markers necessary to obtain a predetermined probability level in detecting non-self-alleles of a given frequency. For instance, the equations can be helpful in the design of assays, where the non-self-allele can be detected by analysis of cfDNA in plasma from pregnant women, to estimate fetal fraction or to monitor changes in cfDNA in plasma of transplantation patients.

The equations give exact, quantitative descriptions of all non-self-situations in pregnancy and transfusion/transplantation.

## Introduction

In immunology, the conceptual dichotomy of self versus non-self underlies the basic tenet that the immune system normally does not react against the individual harboring the immune system, only against antigens alien to the individual. This simple conceptual dichotomy may not fully capture the underlying biological complexity, yet the concept can be useful in immunology. Relatively rare situations in which the immune system reacts against self-components and causes disease are deemed autoimmunity.

Based on this conceptual dichotomy, we wanted to give an exact mathematical description of self versus non-self situations at the population level. A non-self-situation arises when an individual is exposed to an antigen that this individual does not possess. This situation is necessary for the immune system to process and present non-self antigens on e.g. HLA molecules to immune cells. This presentation enables the immune system to launch an allo-immune response. Variations in the primary DNA sequence of exons and silencing of expression may result in different antigen repertoires in other individuals.

A non-self situation can arise in pregnancy when the mother is exposed to an antigen that the fetus has inherited from its father or in recipients of blood transfusion or organ transplantation from an allogeneic donor. Immune responses of the mother against the fetus may lead to injury of the fetus or an immune response against transfused red blood cells may lead to hemolysis and secondary kidney injury. An immune response against a transplanted organ may lead to rejection and death of the transplanted organ.

Simple mathematics was used to describe, in biallelic systems, the fraction of all situations with a non-self-allele irrespective of allele frequency. The non-self-scenarios in both pregnancy and blood donation/transplantation were addressed. The equations presented define the theoretically maximal fraction of situations where an immune response may arise and define all situations where the non-self-allele can be detected by various assays based on primary DNA sequence. The equations presented can be helpful in assay development of non-self in biallelic systems and provide general insight. Most blood group antigens are biallelic as is also much other genetic variation.

Methods for determining fetal fraction have been published, e.g., (Ni et al., 2019).

## Materials and methods

Only biallelic systems with alleles A and B with allele frequencies p and q respectively were considered. Basic mathematics and theoretical deliberations led to the development of three equations describing all non-self-situations in any biallelic system in pregnancy and transfusion/transplantation and based on the equations, simple equations were derived for calculating the number of biallelic markers needed to be combined into an assay to reach a given probability of detecting non-self in pregnancy and transfusion/transplantation including the scenario where both donor and recipient are homozygous albeit with different alleles.

Microsoft Excel was used to evaluate the equations in simulations *in silico* by creating 4000 samples in Hardy-Weinberg equilibrium but with varying allele frequency p of 0.1, 0.2, 0.3, 0.4, and 0.5 respectively. The 4000 samples with the same allele frequency were duplicated and both identical 4000 samples were randomized separately using the SLUMP function in Excel. A 9-digit number between 0 and 1 was generated without duplicate numbers. The genotypes were coupled to the random number and sorted by the number thus generating a column of randomly sorted genotypes. For the prenatal testing, a column with only a single allele was randomized as the paternal contribution to the fetus. The two columns of the independently randomized 4000 samples – with the same allele frequency - were combined into one set of 4000 outcomes and the number of times, a non-self-situation was created, was counted. This was repeated for a total of 4 times for each allele frequency. The number of non-self-outcomes counted was compared with the number of non-self-outcomes predicted as calculated by equations (3), (9) and (12).

The confidence intervals were calculated in GraphPad Prism 10.1.2.

The calculations have the Hardy-Weinberg principle as the basis. In a biallelic system, the frequency of the genotypes in a population is (p+q)^2^=p^2^+q^2^+2pq=1, when there is Hardy-Weinberg equilibrium (Hardy, 1908; Weinberg, 1908).

## Results

### The pregnancy situation

In situations of non-self in pregnancy, the fetus has inherited an allele from its father that the mother does not have. We here extend the term “non-self” to encompass genetic sequence variants that the pregnant woman or recipient of transfusion or organ donation does not have. Irrespective, cfDNA from both the fetus and the pregnant woman is present in maternal plasma and non-self cfDNA sequences can be used as identification tags for the presence of fetally derived cfDNA in pregnant women. Such considerations would also apply to transplantation situations albeit the donor has contributed two alleles.

In the pregnancy situation, the paternal allele inherited by the fetus may or may not be different from the alleles of the mother. In case the fetus inherits an A allele with frequency p_f_ from the father, the _f_ suffix solely denotes that the allele is inherited from the father, it is indiscernible from other p alleles, one of the four situations creates a non-self outcome (Fig 1).

**Fig 1.**
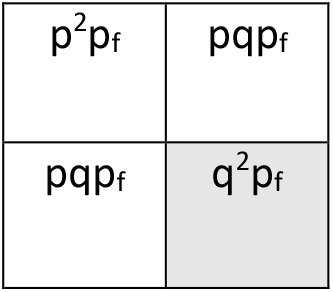
The maternal non-self-situation (cell in grey) in pregnancy is visualized.

If the fetus inherits a B allele with frequency q (q_f_) from the father (Fig 2).

**Fig 2.**
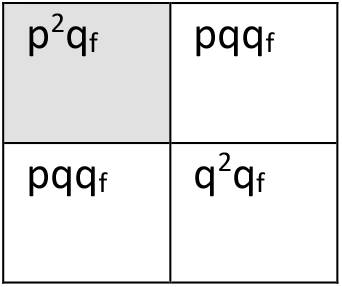
The other maternal non-self-situation (cell in grey) in pregnancy is visualized.

The two situations, p^2^q_f_ and q^2^p_f_ are not the same, but both represent a non-self-situation that can provide the necessary, but not the sufficient basis for immunization and both situations can be informative as to the presence of fetal cfDNA in maternal plasma, i.e., fetal cfDNA that is qualitatively different from the maternal corresponding primary DNA sequence.

This means that for non-self-situations both the p^2^q_f_ and the q^2^p_f_ outcomes are relevant. Mathematically, given that p+q=1, the first situation translates to

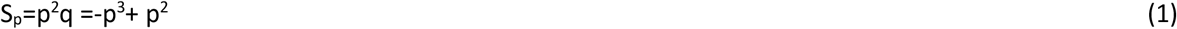

This formula gives the fraction of pregnancies for a given biallelic system where one allele has the frequency p where non-self is present, and the genetic precondition is fulfilled for a possible immunization event (Fig 3).

**Fig 3.**
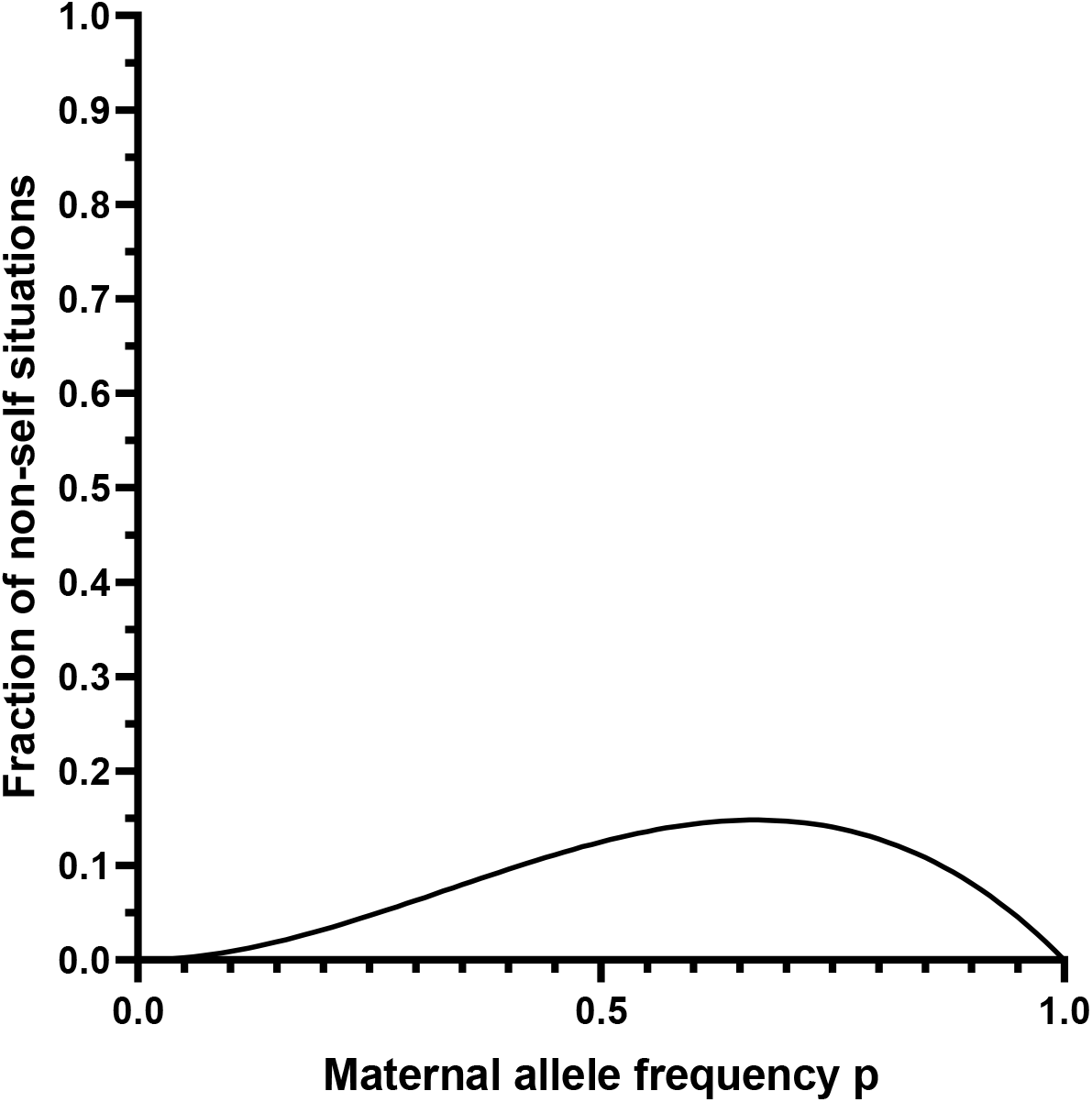
The fraction of pregnancies with non-self of a single allele in a biallelic system. The pregnant woman is p^2^ and the man contributes an allele with frequency q to give a situation with p^2^q described by equation (1): -p^3^+p^2^ (full line).

By integrating equation (1) and calculating the area under the graph in Fig 3, 1/12 of all pregnancies for all values of p have this genetic constellation that is a precondition for maternal immunization for a given single biallelic antigen system. The maximal pregnancy risk allele frequency is found when p is close to 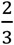 and the risk is low both when p is either very low or very high.

But p^2^q only describes half the possible situations. The formula for the converse situation:

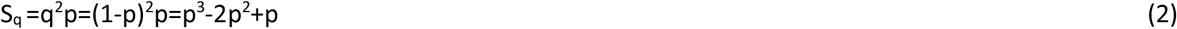

is depicted with a dashed line in Fig 4. With this distribution, the maximal risk situations are when p is close to 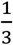 .

**Fig 4.**
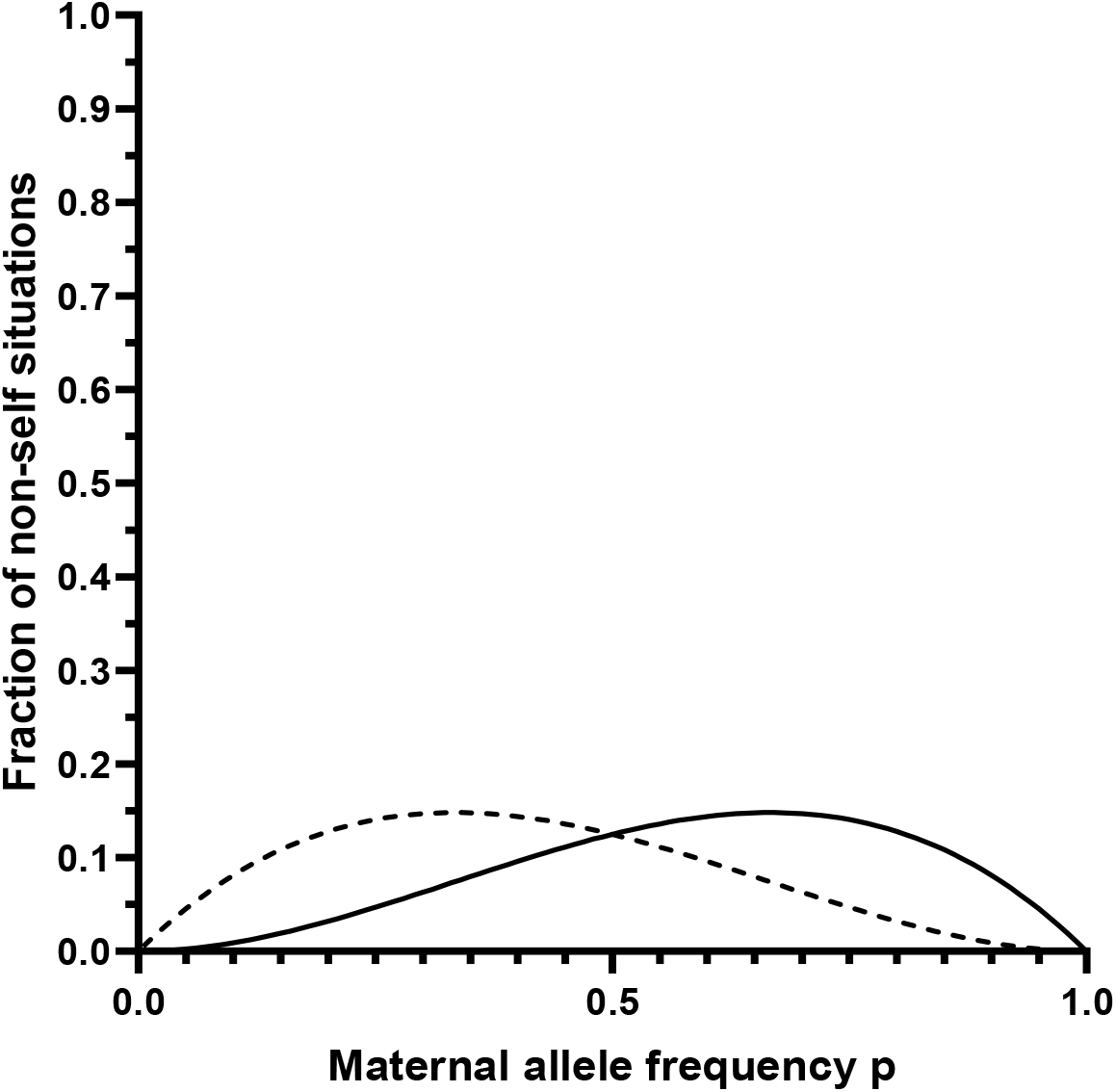
The fraction of non-self in pregnancy in relation to allele frequency for a single biallelic system with the mother being either homozygous p^2^ and the father contributing the other allele with frequency q, giving the p^2^q situation (full line) or homozygous q^2^ with the father contributing an allele with the frequency p, giving the q^2^p situation (dashed line).

The sum of the two distributions is decisive for the total, theoretical immunization risk in a population for a biallelic locus and for all possible non-self-situations in a population. Determining which distribution, a given allele belongs to in a biallelic system is not possible.

In the case of an assay for determining either allele, both equation (1) and (2) must be considered. To obtain a high probability for detection of fetal cfDNA in maternal plasma, it is necessary to use several biallelic variant markers to produce an assay to ascertain if one (or more) alleles, different from the maternal alleles, have been inherited by the fetus. Such an assay will have the added advantage that no prior knowledge of maternal or paternal alleles is needed. An assay addressing this can be useful as a control assay for the presence of fetal cfDNA in maternal plasma when making genetic predictions based on findings of fetal cfDNA in maternal plasma. This principle is not new (Scheffer et al., 2011).

The cumulated information from more marker alleles will help with increased likelihood to establish the presence of fetal cfDNA. Establishing whether fetal cfDNA is present or not will minimize the risk of a false negative result in relation to other tests based on the detection of specific fetal cfDNA. If the presence of fetal cfDNA cannot be ascertained, then a negative result from the detection of specific fetal cfDNA may not be reliable. The principle is illustrated in Fig 5 with an example of five different primer sets targeting five different markers but only the markers on chromosomes 1 and 2 are informative of the presence of fetal cfDNA. So, for one marker either allele may be informative in these situations with both the (p^2^q) and the (q^2^p) situations being informative of the presence of fetal cfDNA in maternal plasma.

**Fig 5.**
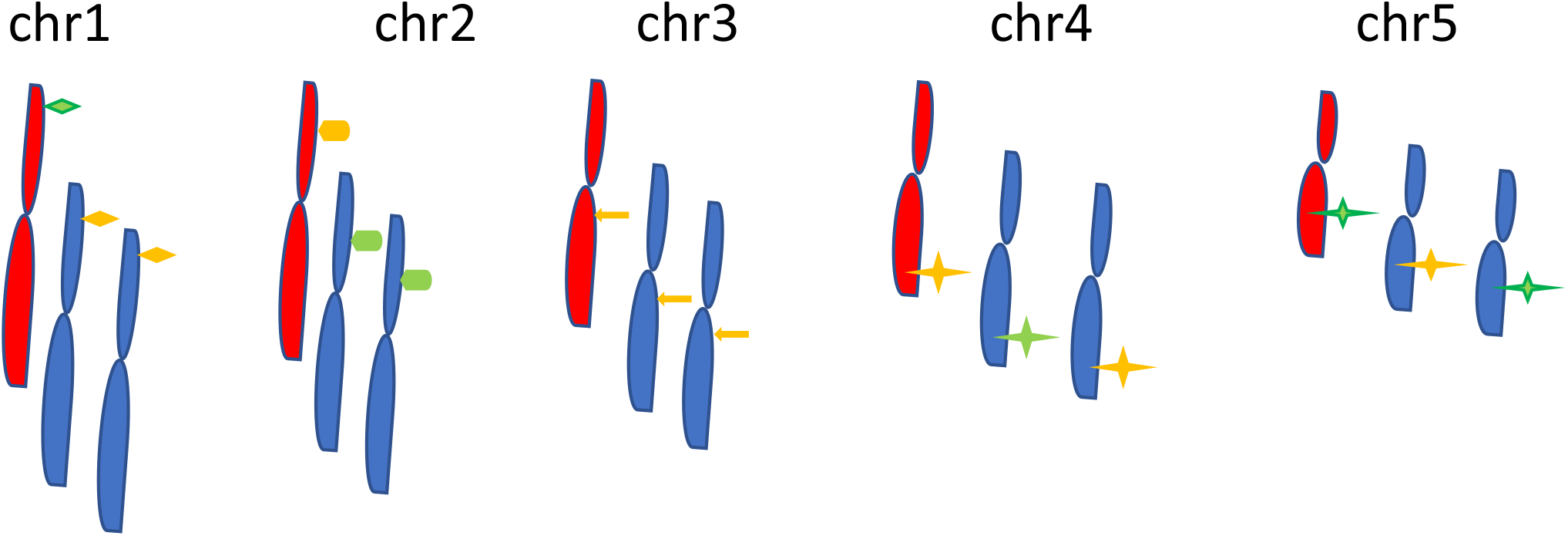
Marker informativity and non-informativity exemplified, (fetal chromosomes red, maternal chromosomes blue).

Both the p^2^q_f_ and the q^2^p_f_ outcome (1/4 of all outcomes from Punnett squares, grey cells) are relevant (Fig 1 and 2) to detect the presence of non-self cfDNA derived from the fetus in maternal plasma. Both situations are theoretically informative (S_I_) of non-self and with risk of immunization of the pregnant woman, and all other situations are non-informative of the presence of fetal cfDNA in maternal plasma, and without risk of immunization of the pregnant woman. The equation for the presence of non-self in the pregnant women will be:

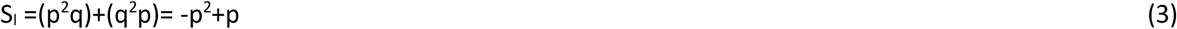

This is shown in Fig 6 for allele frequencies of 0≤p≤1.

**Fig 6.**
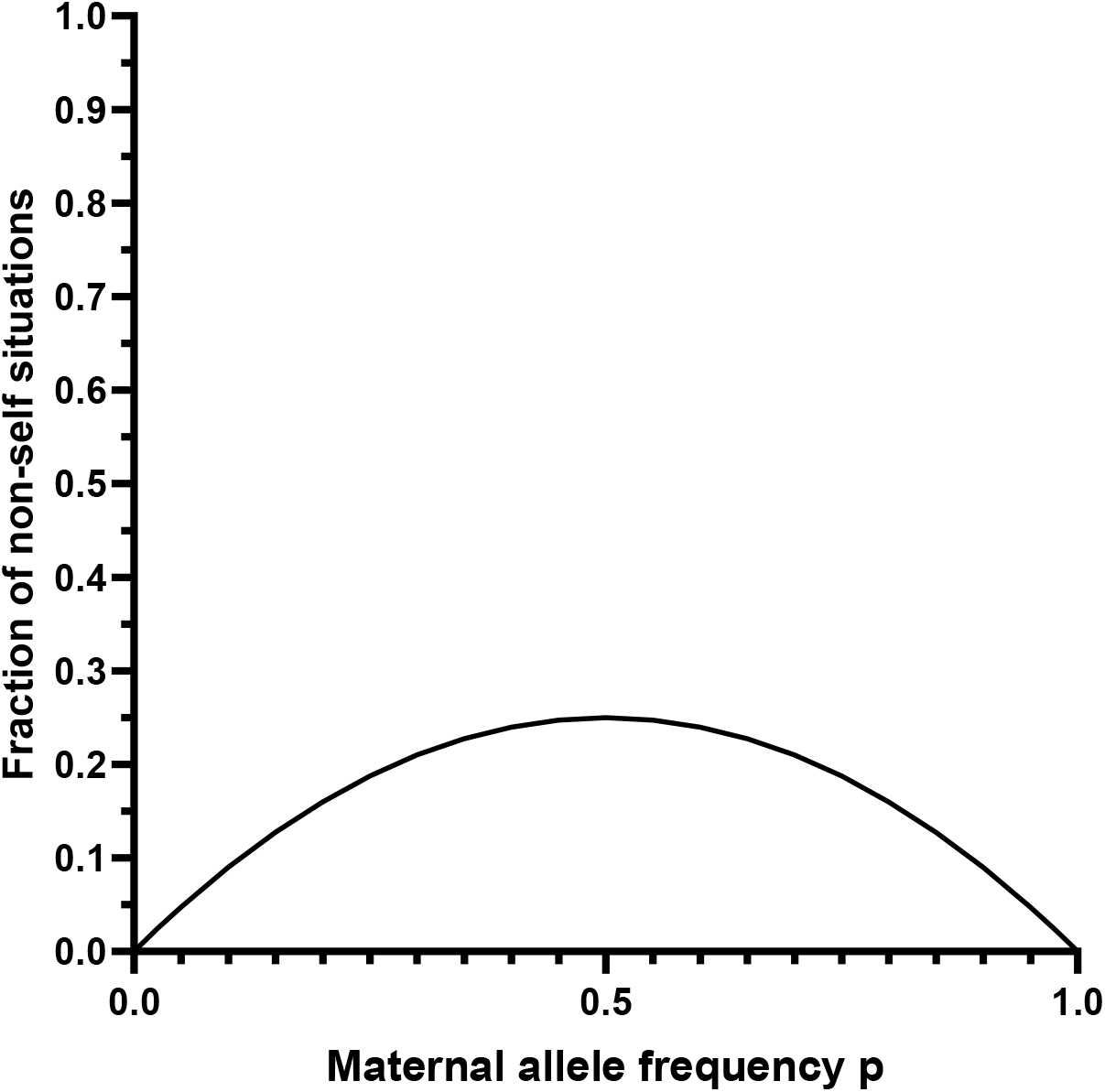
The fraction of all combined non-self outcomes in pregnancy when both p^2^q and q^2^p are included, shown in relation to allele frequency p.

Equation (3) has a maximum at p=0.5. Thus, alleles with a frequency p=0.5 in a biallelic system are optimal for the detection of the presence of fetal cfDNA, as this allele frequency is most informative. By integrating equation (3) and calculating the area under the graph in Fig 6, maximally 1/6 of all situations can be informative in a single biallelic antigen system or result in immunization by contributing a non-self-antigen when both alleles from a biallelic system are taken into consideration. By using alleles with the most informative frequencies in the narrow interval between 0.4 to 0.6, potentially 0.0493 can be accessed, that is 0.0493/0.1667 ∼30% of all theoretically possible information in this setting.

By testing 20 alleles with p=0.5, on average about 5 alleles (0.25×20≈5) will be expected to be non-self and thus informative for a given blood sample; and at least one allele must be informative for an assay to be of use.

A non-informative situation, S_N_ can be calculated by

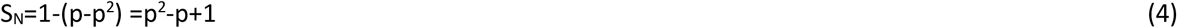

For several alleles with differing allele frequencies p_1_, p_2_, p_3_ …p_i_ an assay, S_I(1-i)_ using these allelic markers will be informative for at least one allelic marker, using equation (4) when

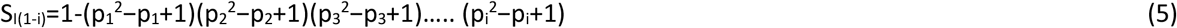

If all allele frequencies are identical p, then equation (5) can be generalized and the fraction of informative situations testing n alleles can be calculated by

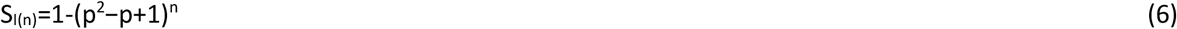

Equation (6) can be rewritten to calculate the number of different allelic markers n, with the same allele frequency p, needed to obtain a desired level of informativity S_I(n)_:

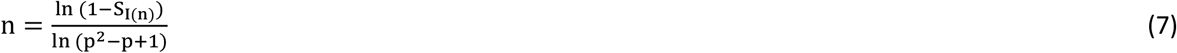

For instance, if information on the presence of fetal cfDNA is wanted in 99% of situations of testing a maternal plasma sample using alleles with a frequency of 0.5, it can be calculated that at least 16 biallelic markers would be needed in an assay as estimated from equation (7). Using for instance 16 markers with an allele frequency of 0.5 would result in 0.9899 probability of detecting a fetal-specific allotype and using 16 markers with an allele frequency of 0.4 would result in 0.9876 of detecting a fetal allotype (equation 6). This is useful information when designing an assay for the detection of fetal cfDNA.

In Fig 7 the application of equation (7) shows the effect of the number of markers with different allele frequencies in relation to the probability of detecting a fetal-specific allotype. Alleles with allele frequencies down to about 0.3 are highly informative and can be included in an assay.

**Fig 7.**
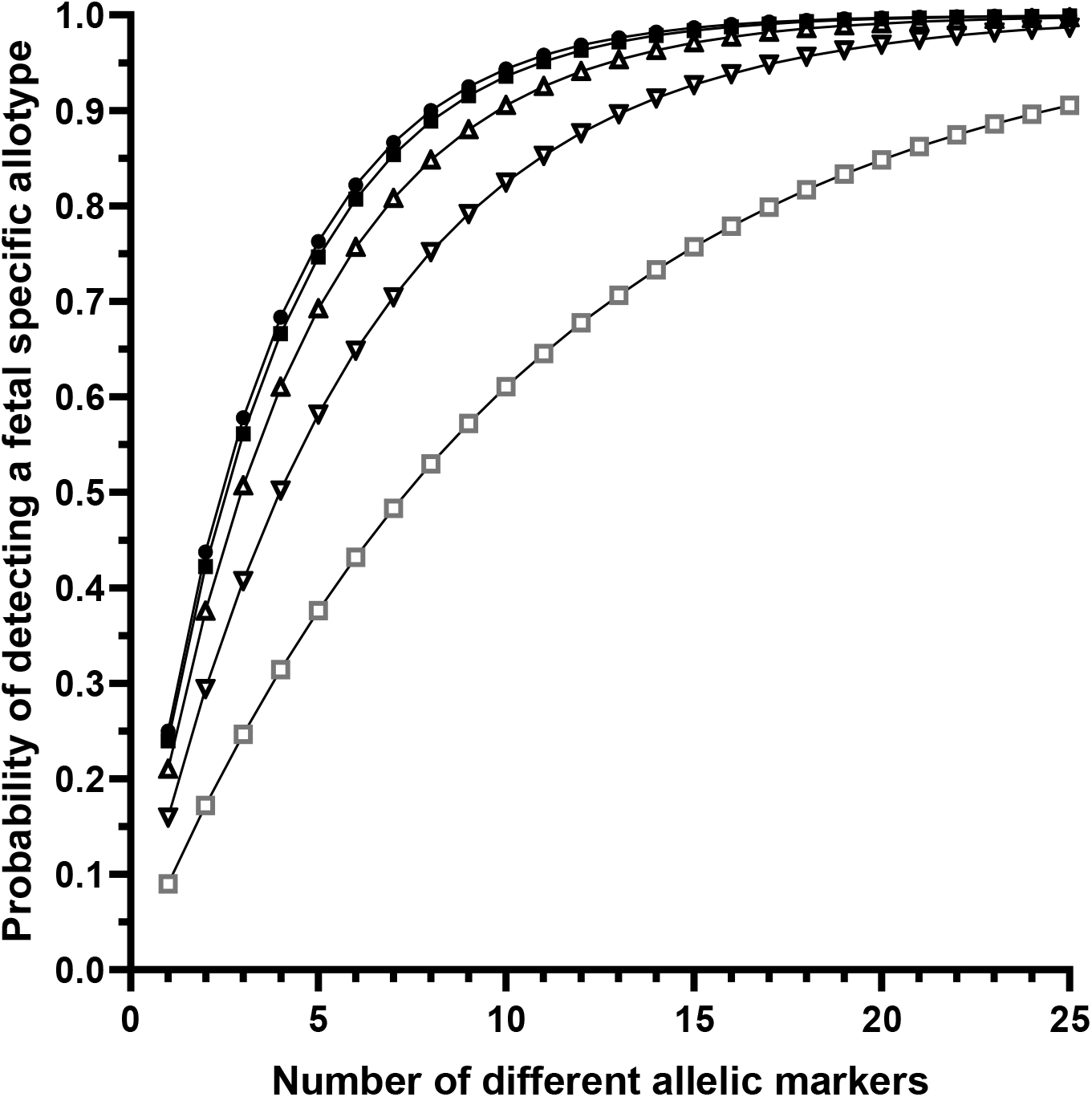
The cumulative effect of choosing multiple biallelic markers for the detection of the presence of non-self in pregnancy i.e., fetal cfDNA in relation to allele frequencies of 0.5 (•), 0.4 (▪), 0.3 (Δ), 0.2 (∇), and 0.1 (□), respectively.

It should be added that if p is substituted with (1-q) in equation (3) -p^2^+p, as –(1-q)^2^+(1-q), the result is -q^2^+q.

An overview of all possible outcomes in pregnancy in relation to allele composition when considering the two maternal alleles and the allele contributed by the father is shown in Fig 8. As the mother will invariably contribute one allele to the fetus (except in situations as the recipient of an egg donation, which situation will be the same as described for the transfusion/transplantation situation) only three alleles are considered. All situations described by equation (3), -p^2^+p give rise to non-self and consecutively risk of immunization as well as being informative in prenatal assays that detect fetal-specific cfDNA sequences. No other situation gives rise to a non-self-situation for the mother.

**Fig 8.**
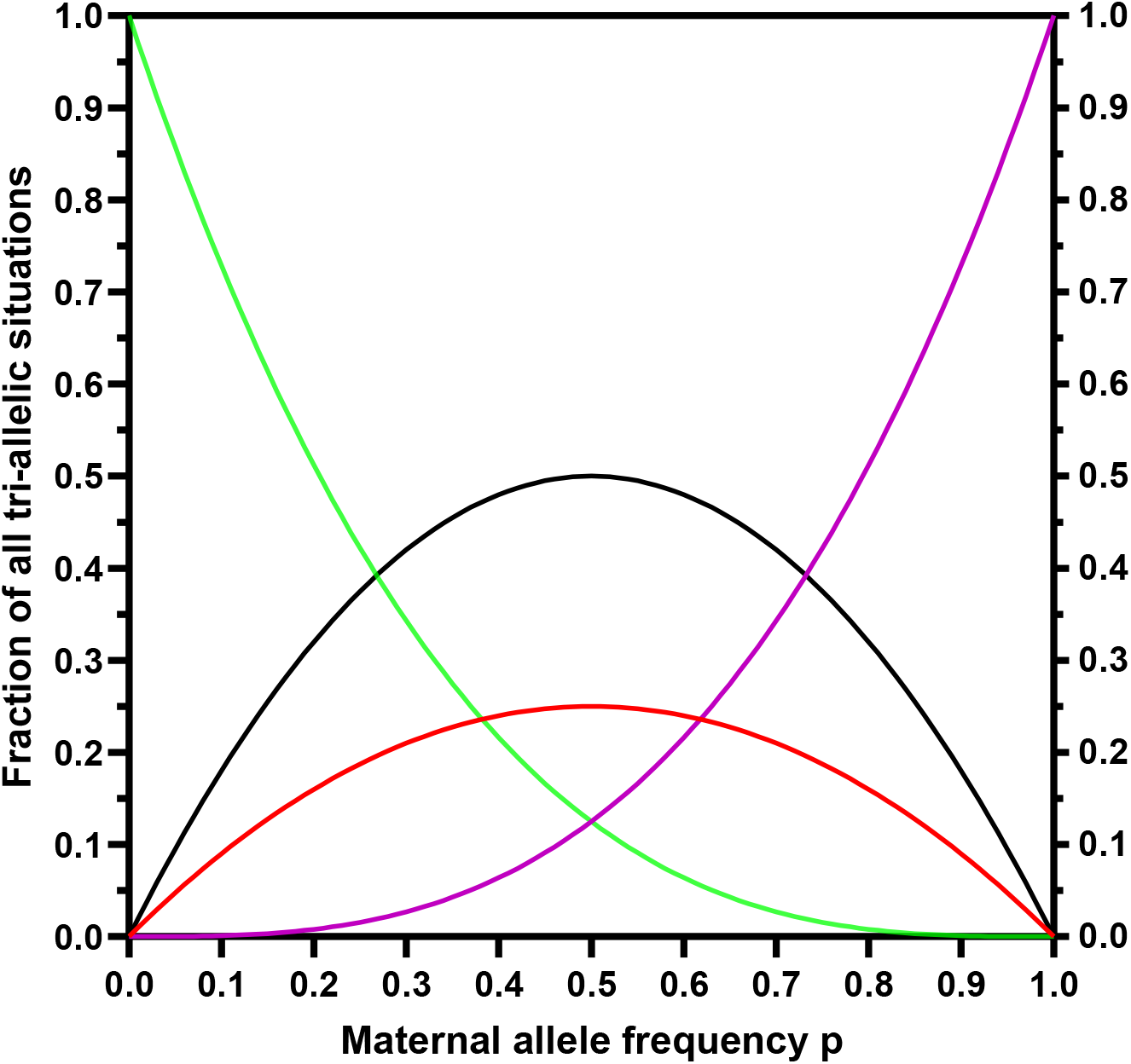
All theoretically possible outcomes of allele combinations in pregnancy for a single biallelic marker of any frequency: p^3^ defined by (p^3^) (lilac graph), q^3^ defined by (-p^3^+3p^2^-3p+1) (green), and the situation with non-self defined by (-p^2^+p) (red), mother heterozygous and fetus with either allele (-2p^2^+2p) (black). All equations added give 1. (-2p^2^+2p-p^2^+p-p^3^+3p^2^-3p+1+ p^3^=1)

If for instance, the allele frequency of p is 0.5, then 25% of all pregnancies will be at risk of immunization, in 12,5% of all pregnancies, the mother is homozygous p^2^ and the fetus has received an allele with frequency q from its father, and in 12,5% of all pregnancies the mother will be homozygous q^2^ and the father has passed on an allele with the frequency p to the fetus. At p=0.5, the mother is heterozygous in 50% of all pregnancies, and in these situations, there is no risk of immunization by either of the alleles.

### The situation of transfusion or transplantation

In the recipient of transfusion/transplantation, non-self-alleles are introduced in the following 4 different recipient-donor situations with the indicated allele frequencies:

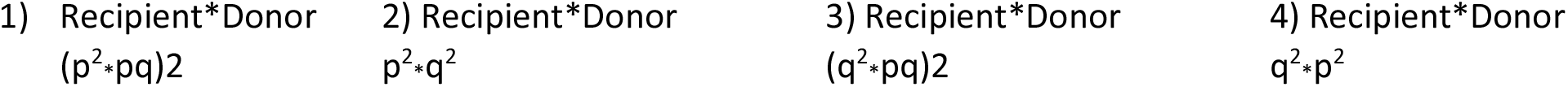

This is shown in Table 1, with cells marked with (#) indicating all the non-self-outcomes. All the non-self-outcomes can be written as:

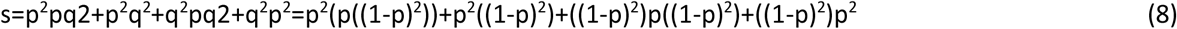

Equation (8) can be simplified as:

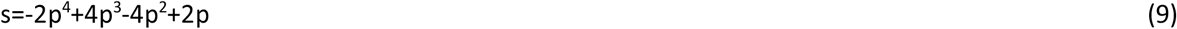

Where p is allele frequency.

**Table 1.** Overview of outcomes in transfusion and transplantation, # and $ marks all non-self-situations for transfusion/transplant recipients.

| donor\recipient | p<sup>2</sup> | q<sup>2</sup> | pq | pq |
| --- | --- | --- | --- | --- |
| p<sup>2</sup> | p<sup>2</sup>p<sup>2</sup> | q<sup>2</sup>p<sup>2</sup> (#,\$) | pqp<sup>2</sup> | pqp<sup>2</sup> |
| q<sup>2</sup> | p<sup>2</sup>q<sup>2</sup> (#,\$) | q<sup>2</sup>q<sup>2</sup> | pqq<sup>2</sup> | pqq<sup>2</sup> |
| pq | p<sup>2</sup>pq (#) | q<sup>2</sup>pq (#) | pqpq | pqpq |
| pq | p<sup>2</sup>pq (#) | q<sup>2</sup>pq (#) | pqpq | pqpq |

And for calculation of n markers with an allele frequency of p, cfr. the equation development for the pregnancy situation:

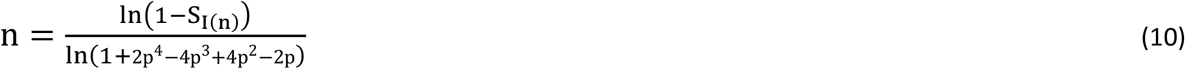

Setting S_I(n)_ at 0.99 and p=0.5 gives n≈10.

This equation can be useful for calculating the number of primer sets needed in design for the detection of non-self in this situation where both a heterozygous and homozygous contribution to non-self can be used for assay purposes.

In Table 1, the cells marked (#) define the non-self-situations that must be considered for assay design when all non-self-situations in transfusion or transplantation must be considered. These situations are described as -2p^4^+4p^3^-4p^2^+2p (equation (9)).

The cells marked ($) define the non-self-situations that must be considered when only the two homozygous situations are relevant e.g., in some assays detecting rejection of a transplanted organ. These situations are described as 2p^4^-4p^3^+2p^2^ (equation (12)).

At p=0.5, 37.5% of recipients of blood transfusion or a recipient of a donor organ will have a non-self-allele for a given biallelic system, equation (9) and Fig 11. An overview of all the fractions in transfusion and organ donation is shown in Fig 9.

**Fig 9.**
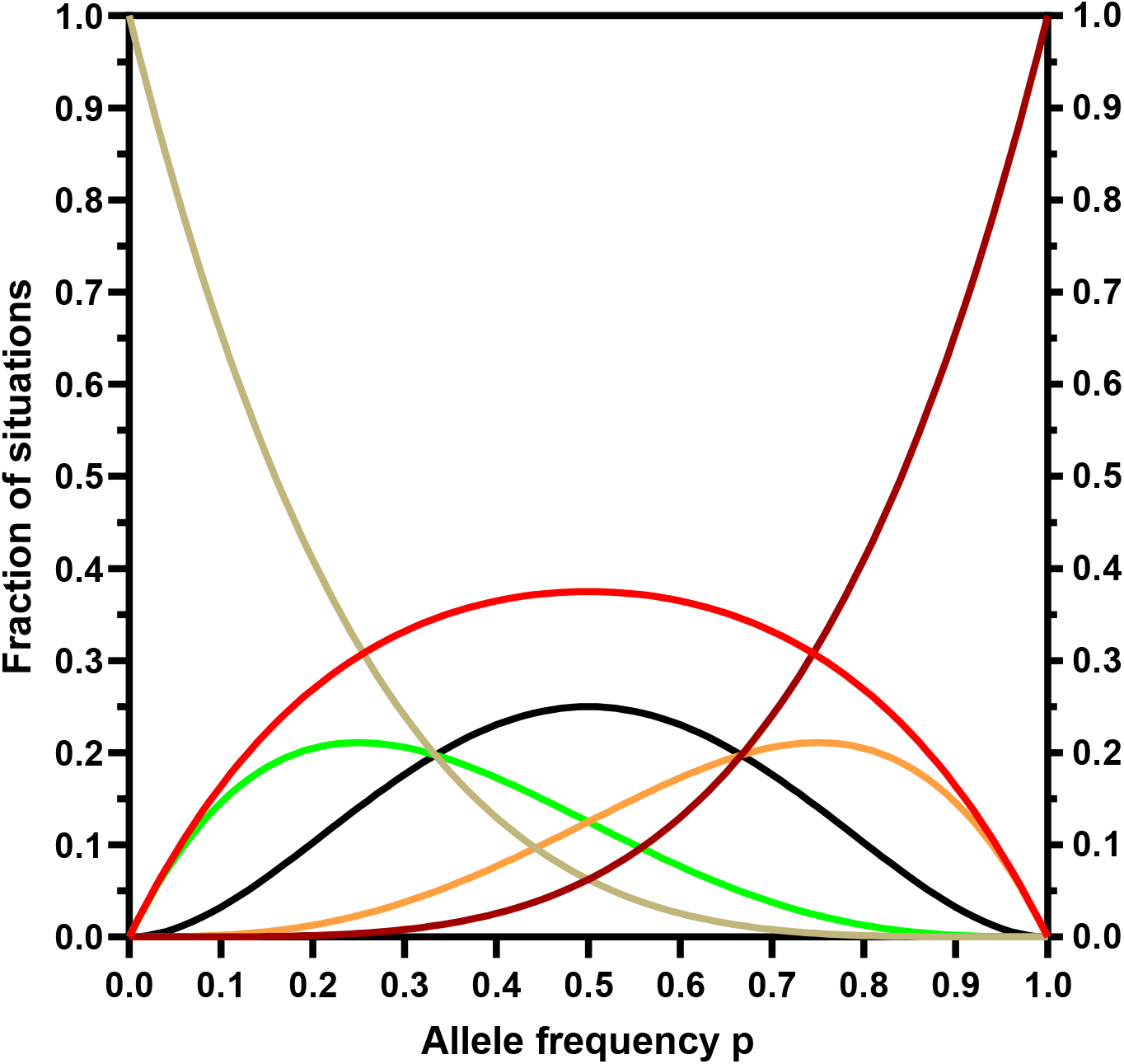
All theoretical situations from transfusion/transplantation in a biallelic system. Situations with non-self are defined by (-2p^4^+4p^3^-4p^2^+2p) (red) and are the combined risk of the two homozygous situations p^2^q^2^ and q^2^p^2^ and the four situations of non-self for homozygous recipient and heterozygous donor p^2^pq+p^2^pq and q^2^pq+q^2^pq (see Table 1). The two latter situations occur with the same fraction as the green and orange graphs respectively. The situations without non-self: with a heterozygous recipient and two different homozygous donors: pqp^2^+pqp^2^ are defined by (-2p^4^+2p^3^) (orange) and pqq^2^+pqq^2^ are defined by (-2p^4^+6p^3^-6p^2^+2p) (green). The situations where the recipient and donor have the same homozygous alleles are defined by p^4^ (p^4^) (brown), and for q^4^ by (p^4^-4p^3^+6p^2^-4p+1) (grey). The 4 situations where both recipient and donor are heterozygous are defined by (4p^4^−8p^3^+4p^2^) (black). All the above equations added give 1. (-2p^4^+4p^3^-4p^2^+2p+p^4^+p^4^-4p^3^+6p^2^-4p+1-2p^4^+2p^3^-2p^4^+6p^3^-6p^2^+2p+4p^4^−8p^3^+4p^2^=1)

**Fig 10.**
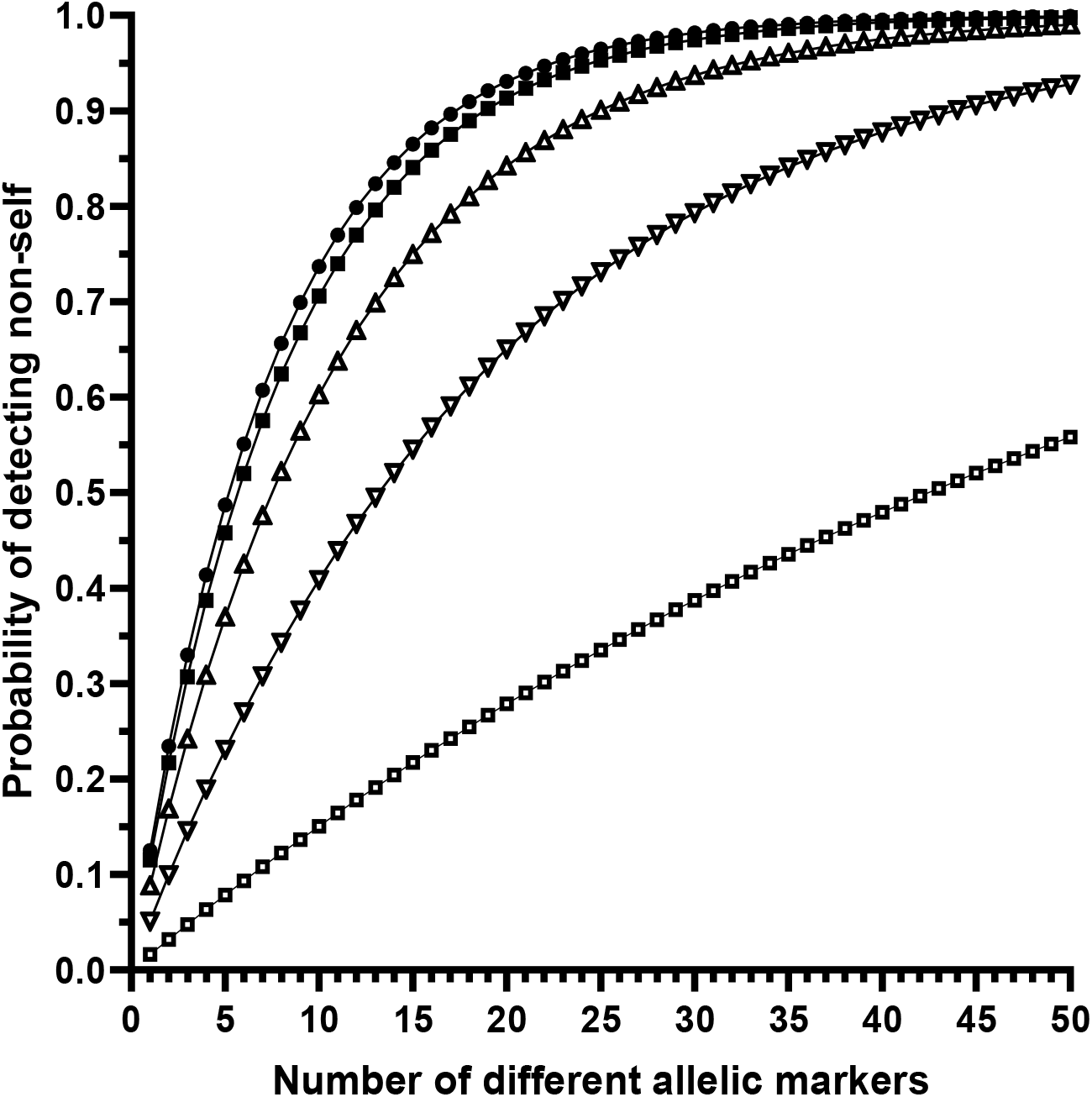
The probability of detecting non-self in relation to allele frequencies and the cumulative number of markers used in an assay of the double homozygous situation. Allele frequencies of 0.5 (•), 0.4 (▪), 0.3 (Δ), 0.2 (∇), and 0.1 (□), respectively are shown.

**Fig 11.**
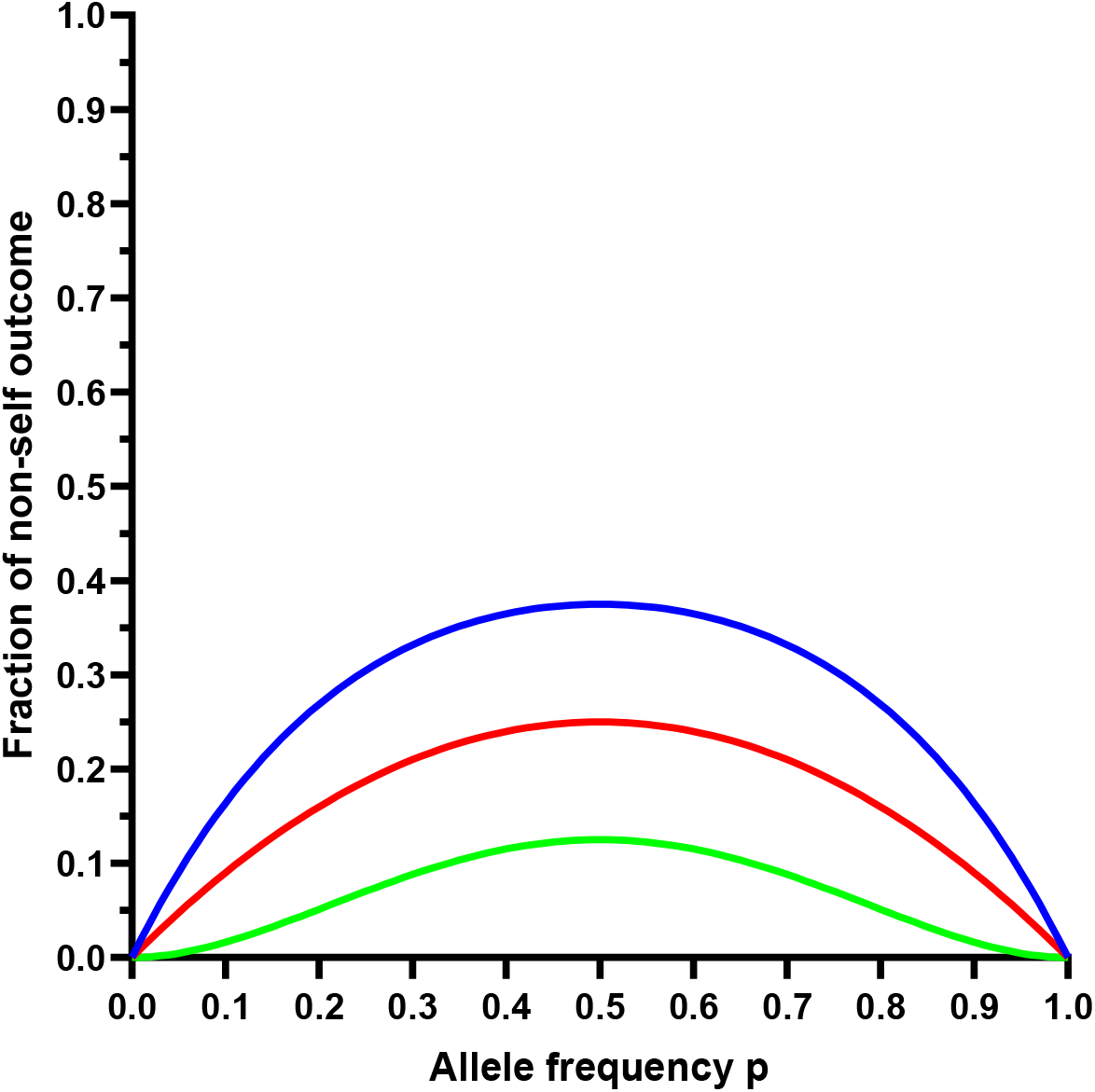
The fraction of non-self-outcomes predicted by the three equations as a function of allele frequency. The double homozygous non-self-situations defined by (2p^4^-4p^3^+2p^2^) (green), the non-self-situations relevant for pregnancy defined by (-p^2^+p) (red), and all non-self-situations in transfusion and transplantation are defined by (-2p^4^+4p^3^-4p^2^+2p) (blue).

From Fig 9 it can be seen that non-self-situations (the red graph) arise at a fairly constant level for values of p between 0.2 to 0.8 and make up a fraction of about 0.25-0.35 in this interval of p. None of the other genotype combinations pose any risk as to immunization of the recipient. The two equations for single alleles in pregnancy have maximal fractions exactly where the three graphs intersect at p=1/3 and 2/3 respectively.

In other situations, where it is desirable to detect admixed DNA from different individuals such as for chimerism measurements in HSCT or some cases of organ transplantation it can be desirable to investigate only the double homozygous situation; the mathematics is slightly different.

An optimally informative situation S_I_ for e. g. digital PCR to monitor cfDNA in cases of chimerism for instance after transplantation would be when the donor is homozygous for a given marker and the recipient is homozygous for the other allele or vice versa (cells marked ($) in table 1):

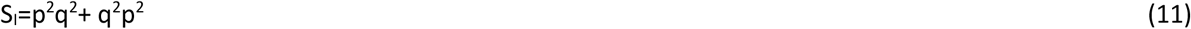

Given that p+q=1, this can be simplified to

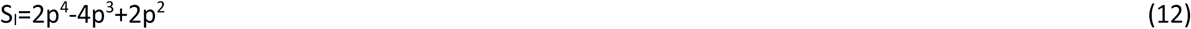

And the situation F_N_ that is non-informative

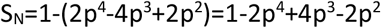

For more markers with varying allele frequencies p_1_, p_2_, p_3_…p_i,_ where at least one situation is informative.

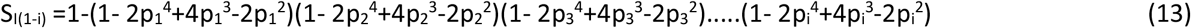

When p is the same for n different allelic markers, the formula can be simplified to

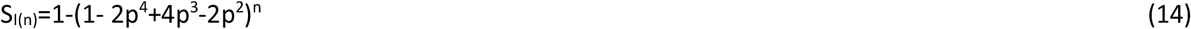

Rearranged:

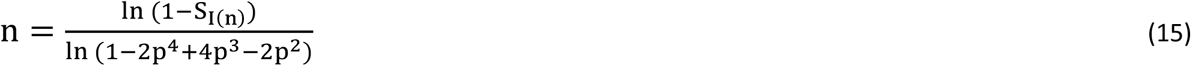

Setting S_I(n)_ at 0.99 and p=0.5 gives n≈34, i.e., 34 primer sets with biallelic markers with an allele frequency of p=0.5 are needed to have, with 99% probability, at least one marker that can be used to discern between recipient and donor cells with a marker that is homozygous in the recipient as well as homozygous in the donor albeit for the other allele. If p=0.4 then 38 allelic markers would be needed to achieve the same S_I(n)_. With p=0.5 a check of equation (15) for 34 markers gives S_I(n)_= (1-(2/16))^34^=0.0107

In Fig 10 the application of equation (14) shows the number of markers with different allele frequencies needed to obtain a given level of probability of detecting non-self. Alleles with allele frequencies down to about 0.4 are highly informative and can be included in an assay.

The graphs in Fig 11 depict the 3 different scenarios described by the three different equations for the contribution of non-self. At p=0.5, the graph describing the transfusion/transplantation situation has a maximum of 6/16, the graph describing pregnancy has a maximum of 4/16, and the graph describing the double homozygous situation has a maximum of 2/16, corresponding to the fractional number of cells in Table 1 with the genotypes used for the deduction of the equations.

The result of the simulation of the combined 2×4000 constructed and randomized genotypes in Hardy-Weinberg equilibrium showed good agreement with the results predicted by the equations (Fig 12) and Table 2. The number of non-self-situations that were counted from the simulation of the transfusion/organ recipient situation, the pregnancy situation, and the double homozygous scenario, were compared to the predicted numbers from the equations (9), (3), and (12). There was no significant difference in Fisher’s exact test at p<0.05.

**Table 2.** The numbers from the simulations and predictions with an allele frequency of p that underlies Fig 12 are shown. Found_tx and predicted_tx refer to transplantation/transfusion situations, found_pre and predicted _pre refer to the prenatal situation, and found_hom and predicted_hom refer to the double homozygous situation.

| Allele frequency | found_tx | predicted_tx | found_pre | predicted_pre | found_hom | predicted_hom |
| --- | --- | --- | --- | --- | --- | --- |
| 0.1 | 659 | 655.2 | 357 | 360 | 64 | 64.8 |
|  | 644 |  | 357 |  | 68 |  |
|  | 669 |  | 362 |  | 64 |  |
|  | 637 |  | 344 |  | 59 |  |
| 0.2 | 1068 | 1075.2 | 629 | 640 | 200 | 204.8 |
|  | 1089 |  | 660 |  | 202 |  |
|  | 1056 |  | 628 |  | 193 |  |
|  | 1044 |  | 632 |  | 200 |  |
| 0.3 | 1324 | 1327.2 | 840 | 840 | 351 | 352.8 |
|  | 1313 |  | 865 |  | 328 |  |
|  | 1324 |  | 837 |  | 356 |  |
|  | 1344 |  | 824 |  | 343 |  |
| 0.4 | 1493 | 1459.2 | 980 | 960 | 458 | 460.8 |
|  | 1454 |  | 985 |  | 462 |  |
|  | 1448 |  | 971 |  | 442 |  |
|  | 1444 |  | 982 |  | 437 |  |
| 0.5 | 1512 | 1500 | 998 | 1000 | 519 | 500 |
|  | 1510 |  | 980 |  | 492 |  |
|  | 1508 |  | 996 |  | 509 |  |
|  | 1456 |  | 1030 |  | 490 |  |

**Fig 12.**
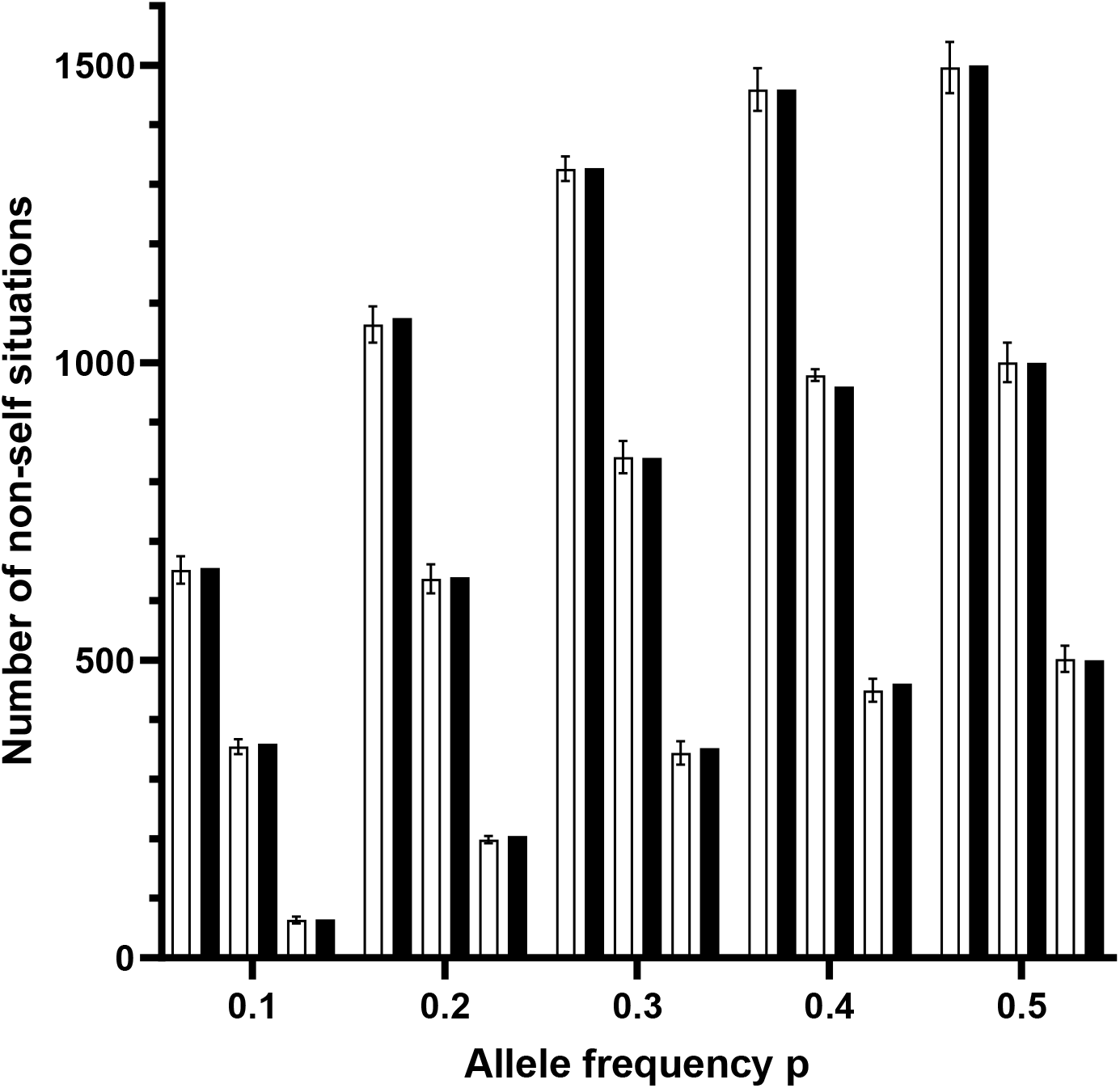
*In silico* simulation. Non-self-situations for transplantation, pregnancy, and double homozygous scenarios were counted after *in silico* simulation of 4000 constructed alleles/genotypes in Hardy-Weinberg equilibrium with 5 different allele frequencies (0.1, 0.2, 0.3, 0.4, and 0.5), each with four replicates. The counted non-self-situations (white columns) were compared to the predicted situations (black columns) by the equations (9), (3), and (12). The 95% confidence interval is shown for the counted situations. For each allele frequency, the two first columns are from the transfusion/transplantation scenario, the next two columns are from the pregnancy scenario and the last two columns are from the double homozygote scenario.

The biggest differences were found at allele frequency of 0.4 for the prenatal simulation with a mean of 980 and a predicted number of 960.

The 95% confidence intervals were calculated based on four generated replicates of random genotype combinations and all except one predicted value (p=0.4 for pregnancy) were within the 95% confidence intervals.

The simulation was also done once with 400 samples with consistent results (data not shown).

The three polynomial equations (9), (3), and (12) can be characterized for all theoretical outcomes valid for 0≤p≤1 and the fraction ≥0 and ≤1:

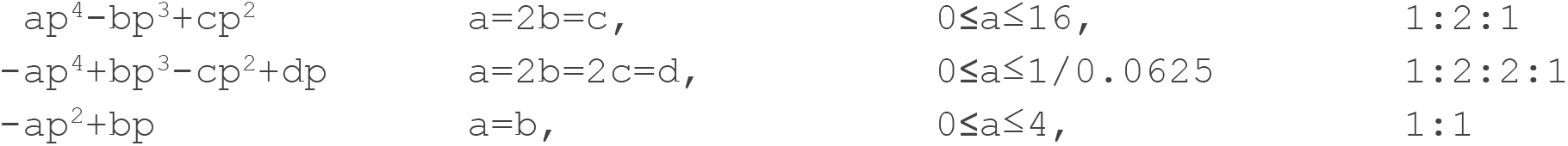

However, only equations (9), (3), and (12), are valid in the stated biological context.

## Discussion

The premise for the mathematical description of the number of markers needed is that alleles have undergone random assortment in accordance with Mendel’s third law which says that alleles are independently assorted and thus that traits encoded by the alleles segregate independently of each other during gamete formation. The physiologic process of meiosis with independent segregation of non-disequilibrium alleles thus underlies the mathematical descriptions. In general, the same assumptions that apply to Hardy-Weinberg calculations would apply to the mathematical description.

Different equations describe the fraction of non-self in pregnancy on one hand and transfusion and transplantation on the other hand, which makes biological sense as in pregnancy, the father always passes only one of his two alleles to the fetus. The two scenarios of transfusion and transplantation are analogous in respect to the description of non-self.

All the non-self-scenarios in biallelic systems were described quantitatively for all values of p in pregnancy, transfusion/transplantation, and the special situation of a homozygous recipient and a donor homozygous for the other allele. In these situations, easy calculation of the number of markers needed for obtaining a desired information level in assays can be obtained regarding the presence of non-self-genetic variants from equations (7), (10), and (15).

Non-self-situations are the prerequisite for an alloimmune response, although several additional conditions must be fulfilled for an immune response to occur.

Assays for determining chimerism in transplantation without prior knowledge of the genotypes of the involved individuals have been developed (Clausen et al., 2023). For instance, one group has chosen 24 indel markers using both homozygous and heterozygous informative marker genotypes (Pettersson et al., 2021).

The equations can be used to assess the number of markers to be used in prenatal control assay for the presence of fetal cfDNA to minimize the risk of false negative results. Also, it is important to note that the graph describing the pregnancy situation has a form that indicates that alleles with allele frequencies far from the optimal 0.5 are very informative as to non-self (Fig 6). This is also indicated in Fig 7 where alleles with a frequency as low as 0.3 appear to be reasonably informative (Lee et al., 2017).

By comparing Fig 7 and Fig 10, clearly, more markers are needed in the double homozygous situation to obtain the same probability of detecting non-self as compared to the pregnant situation.

In the case of the double homozygous situation, a large number of assays with individual markers must be designed to ensure a test with a high rate of useful outcomes. However, the markers should be assorted independently and therefore be spaced sufficiently. With a distance between loci of 40 GB (LaRue et al., 2014) about 75 markers can be designed from the human genome for a single assay.

The equations can also be used to calculate the expected frequency with which a given assay will fail to produce information on presence of fetal cfDNA.

In forensics multiallelic STR systems are routinely used and estimations of the number of biallelic SNPs needed to replace STRs have been done (Amorim and Pereira, 2005; Gill, 2001; Lee et al., 2017).

The mathematical descriptions were tested *in silico* to ascertain that the mathematical predictions were accurate. There was no significant deviation (at p<0.05) by Fisher’s exact test from the counted versus the expected numbers as calculated by equations (9), (3), and (12), see Fig 12. Thus, this *in silico* test does not invalidate the predictive accuracy of the equations, however, a more rigorous *in silico* validation would need both a much larger sample size and many more replicates. In three situations the predicted numbers fell just outside the 95% confidence interval, with a total of 4 replicates. With so many calculations, this is not surprising.

In conclusion, a mathematical description of non-self-allele fractions of biallelic systems is reported in 3 different scenarios: pregnancy, transfusion/transplantation including the scenario with a homozygous donor and a recipient homozygous for the other allele. Also given, are equations to calculate the number of marker systems needed to reach a given probability of detecting non-self. These equations can be useful in quantitative estimations including for the design of tests for identification purposes e.g., fetal fraction or chimerism determination and other purposes.

## Abbreviations

cfDNA: cell-free DNA
GB: gigabase
HLA: Human Leucocyte Antigen
HSCT: Human Stem Cell

## References

Amorim, A. and Pereira, L., 2005. Pros and cons in the use of SNPs in forensic kinship investigation: a comparative analysis with STRs. Forensic Sci Int. 150, 17–21.

Clausen, F.B., Jorgensen, K.M.C.L., Wardil, L.W., Nielsen, L.K. and Krog, G.R., 2023. Droplet digital PCR-based testing for donor-derived cell-free DNA in transplanted patients as noninvasive marker of allograft health: Methodological aspects. PLoS One. 18, e0282332.

Gill, P., 2001. An assessment of the utility of single nucleotide polymorphisms (SNPs) for forensic purposes. Int J Legal Med. 114, 204–10.

Hardy, G.H., 1908. Mendelian proportions in a mixed population. Science. 28 49–50.

LaRue, B.L., Lagace, R., Chang, C.W., Holt, A., Hennessy, L., Ge, J., King, J.L., Chakraborty, R. and Budowle, B., 2014. Characterization of 114 insertion/deletion (INDEL) polymorphisms, and selection for a global INDEL panel for human identification. Leg Med (Tokyo). 16, 26–32.

Lee, H.J., Lee, J.W., Jeong, S.J. and Park, M., 2017. How many single nucleotide polymorphisms (SNPs) are needed to replace short tandem repeats (STRs) in forensic applications? Int J Legal Med. 131, 1203–1210.

Ni, M., Peng, X.L. and Jiang, P., 2019. Bioinformatics Pipeline for Accurate Quantification of Fetal DNA Fraction in Maternal Plasma. Methods Mol Biol. 1909, 177–180.

Pettersson, L., Vezzi, F., Vonlanthen, S., Alwegren, K., Hedrum, A. and Hauzenberger, D., 2021. Development and performance of a next generation sequencing (NGS) assay for monitoring of mixed chimerism. Clin Chim Acta. 512, 40–48.

Scheffer, P.G., de Haas, M. and van der Schoot, C.E., 2011. The controversy about controls for fetal blood group genotyping by cell-free fetal DNA in maternal plasma. Curr Opin Hematol. 18, 467–73.

Weinberg, W., 1908. Über den Nachweis der Vererbung beim Menschen. Jahresh. Ver. Vaterl. Naturkd. 64 369–382.

